# Learning decouples accuracy and reaction time for rapid decisions in a transitive inference task

**DOI:** 10.1101/2025.02.11.636952

**Authors:** Fabian Munoz, Greg Jensen, Maxwell Shinn, Yelda Alkan, John D. Murray, Herbert S. Terrace, Vincent P. Ferrera

## Abstract

Transitive inference (TI) is a cognitive process in which decisions are guided by internal representations of abstract relationships. While the mechanisms underlying transitive learning have been well studied, the dynamics of the decision-making process during learning and inference remain less clearly understood. In this study, we investigated whether a modeling framework traditionally applied to perceptual decision-making—the drift diffusion model (DDM)—can account for performance in a TI transfer task involving rapid decisions that deviate from standard accuracy and response time (RT) patterns. We trained six macaque monkeys on a TI transfer task, in which they learned the implied order of a novel list of seven images in each behavioral session, indicating their decisions with saccadic eye movements or reaching movements. Consistent learning of the list structure was achieved within 200–300 trials per session. Behavioral performance exhibited a symbolic distance effect, with accuracy increasing as the ordinal distance between items grew. Notably, RTs remained relatively stable across learning, despite improvements in accuracy. We applied a generalized DDM implementation (PyDDM; Shinn et al., 2020) to jointly fit accuracy and RT data. Model fits were achieved by incorporating both an increasing evidence accumulation rate and a collapsing decision bound, successfully capturing the RT distribution shapes observed during learning. Learning and transfer were fit by varying drift rate with little change in other parameters. Eye and reaching movements showed similar dynamics, with the difference in RT accounted for mainly by non-decision time. Our results highlight a distinct dynamical regime of the DDM framework, extending its applicability to cognitive domains involving symbolic reasoning and serial relational learning.

## Introduction

Transitive inference (TI) assesses the ability to learn and generalize relationships that obey the principle of transitivity—for example, if A > B and B > C, it follows that A > C. This cognitive ability is typically tested by training subjects on novel lists comprising 5–9 easily discriminable items, such as images, and examining their performance on untrained item pairs. Extensive empirical evidence suggests that TI performance is supported by the formation of an internal representation of the list’s ordinal structure (Prado et al., 2012; Bonnefond et al., 2014; Jensen et al., 2015). However, the mechanisms by which such internal representations are translated into behavioral responses remain poorly understood. In particular, it is unclear whether the decision-making processes underlying TI share properties with those involved in perceptual decision-making.

In the present study, we examined the decision dynamics of nonhuman primates performing a TI task. In each session, monkeys were presented with novel sets of images in paired comparisons and were required to infer their implicit order. While decision accuracy increased with learning, response times (RTs) remained fast and relatively constant, despite the absence of strong time constraints. These findings suggest a dissociation between accuracy improvements and temporal dynamics of decision-making, raising questions about how internal representations are accessed and acted upon during learning.

TI differs fundamentally from traditional perceptual decision-making tasks in that it relies on internally constructed rather than externally available evidence. Consequently, variations in performance are more closely tied to abstract task variables than to sensory stimulus properties. One such variable is symbolic distance (SD)—defined as the difference in rank between two items in the inferred list. Greater symbolic distance is strongly associated with higher accuracy, a hallmark of TI performance. This reliance on internally guided inference suggests that the dynamics of TI decisions may diverge from the patterns observed in perceptual tasks, where decision times typically vary systematically with stimulus strength.

In many two-alternative choice paradigms involving nonhuman primates, RTs often exceed a full second and are inversely correlated with stimulus strength for both dynamic (Roitman & Shadlen, 2002; Churchland et al., 2008; Ding & Gold, 2010) and static stimuli (Ratcliff et al., 2003; Middlebrooks & Schall, 2014). These behavioral metrics are closely linked to the temporal evolution of neural activity across cortical and subcortical regions, including the lateral intraparietal area (Roitman & Shadlen, 2002), prefrontal cortex (Kim & Shadlen, 1999; Kiani et al., 2014), caudate nucleus (Ding & Gold, 2010), and superior colliculus (Jun et al., 2021). These observations have informed the development of computational models, particularly the drift-diffusion model (DDM), which provide principled explanations for the coupling between accuracy and RT in perceptual decisions.

Recent work highlights the existence of decision processes across multiple timescales (Murray et al., 2014; Soltani et al., 2021), including rapid decisions in naturalistic contexts, where speed is critical for survival. Indeed, studies of oculomotor behavior reveal that primates can make highly accurate decisions within 200–300 milliseconds, with frontal eye field (FEF) activity discriminating targets within this timeframe (Cohen et al., 2010; Stanford et al., 2010; Shankar et al., 2011).

The Drift Diffusion Model (DDM)—a widely used framework grounded in sequential sampling theory (Wald, 1945; Ratcliff, 1978)—treats decisions as a process of accumulating noisy evidence until a response threshold is reached. This approach captures key empirical regularities in accuracy and RT across many perceptual tasks. However, its application to tasks such as TI, where performance is not directly tied to sensory input, remains contentious. The qualitative differences between TI and perceptual paradigms motivate a critical examination of whether the DDM can meaningfully capture decision behavior in this domain.

To address this, we applied a generalized drift-diffusion model (GDDM) using PyDDM (Shinn et al., 2020) to simultaneously fit accuracy and RT data from monkeys learning novel TI problems. The GDDM provided high-fidelity fits to both behavioral measures. Model estimates indicated that both the drift rate (reflecting the strength of evidence accumulation) and boundary separation (reflecting response caution) varied as a function of learning, with a sharp increase at the point of transfer. Drift rate was also strongly modulated by symbolic distance. Differences between eye and reaching movements were reflected mainly in non-decision time rather than decision dynamics. These findings contrast with those from other domains, where constant decision boundaries are typically sufficient (e.g., Voskuilen et al., 2017) and suggest that TI behavior is best captured by a collapsing bound model. Our results thus extend the DDM framework to a novel cognitive domain, illuminating the temporal dynamics of decisionmaking based on internal structure rather than perceptual input.

## Materials and Methods

### Subjects

Subjects were six male rhesus macaques. The NHP were 13-14 years old and weighed 8-14 kg at time of experiment. Three animals (F, H, L) performed the task using eye movements as the operant response. Three other NHP (O, Q, R) performed the task by reaching out to a touchpanel display. The NHP that were trained to use touchpanels were never trained to perform TI or any other task using eye movement responses. The research was conducted in accordance with U.S. Department of Health and Human Services (National Institutes of Health) guidelines for the care and use of laboratory animals and was approved by the Institutional Animal Care and Use Committee at Columbia University and the New York State Psychiatric Institute. Monkeys were prepared for experiments by surgical implantation of a post for head restraint and a recording chamber to give access to prefrontal cortex. All surgery was performed under general anesthesia (isoflurane 1-3%) and aseptic conditions. Subjects had prior experience performing transitive inference tasks (Jensen et al. 2015).

The number of animals was chosen to establish reproducibility while avoiding redundancy, and to compare eye movement and reaching responses. For train/test comparisons, each animal served as its own control. In-house software was used to conduct power analyses to determine the number of trials needed to detect changes in accuracy and reaction time. Accuracy (proportion correct) was modeled as a binomial variable. To detect a change from 50% to 60% correct at a power level of 0.8 required 187 trials. Estimates of NHP saccade reaction time variability were taken from a published study (Lawrence et al., 2008). To detect a change in RT from 200 to 210 msec with a standard deviation of 20 msec required a sample size of 34 trials.

### Visual Stimuli and Eye/Reaching Movements

Nonhuman primates (NHPs) were seated in an upright primate chair during the experiment. Three NHPs responded to visual stimuli by making saccadic eye movements. Stimuli were generated and controlled by a CRS VSG2/3F video frame buffer and displayed on a CRT or LCD monitor with a resolution of 1280 x 1024 pixels at 60 Hz. The viewing distance for eye movements was 60 cm. Three other NHP responded with reaching movements to stimuli presented on an LCD panel (1280 x 1024 pixels at 60 Hz) with an integrated resistive touch mechanism (ELO Touch, Rochester, NY). The viewing distance for reaching movements was 25 cm.

Visual stimuli consisted of 140-by-130 pixel color photographs (7° by 8° visual angle for eye movement, 14° by 16° for reaching movements) and small squares that served as fixation and eye movement targets. To control the retinal position of visual stimuli, monkeys were required to maintain fixation before the stimuli were presented. The fixation point was a small red square (0.5° visual angle). NHP were allowed to initiate a saccade response any time after stimulus onset.

Eye position was recorded using a monocular scleral search coil (CNC Engineering, Seattle WA) and digitized at a rate of 1 kHz (Judge et al. 1980). Subjects expressed their choices by making eye movements from the central fixation point to peripheral target stimuli. Eye velocity was computed offline by convolving eye position with a digital filter. The filter was constructed by taking the first derivative of a temporal Gaussian, *G*(*t*), such that 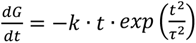, where τ = 8 msec and *k* is a constant that sets the filter gain to 1.0. This filter does not introduce a time shift between the position input and velocity output, but adds temporal uncertainty to the velocity estimates. Horizontal and vertical eye velocities (*h*′(*t*) and *v*′(*t*), respectively) were combined to estimate radial eye speed *r*′(*t*) using the formula 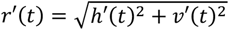. Touch coordinates (position and time) were read directly from the touchpanel software driver in mouse emulation mode.

### Transitive Inference Task

Prior to the beginning of each behavioral session, a set of 7 photographs never before presented to the subjects was selected from a database of over 2500 images. A different set of images was used for each behavioral session. These stimuli were randomly assigned a unique rank, indicated by the letters A thru G, to create an ordered list (**Fig. 1A**). No explicit information about stimulus rank was presented to the subjects. That is, the rank assigned to each stimulus was never displayed, nor was there any information about serial order conveyed by the spatial or temporal presentation of the stimuli.

**Figure 1.**
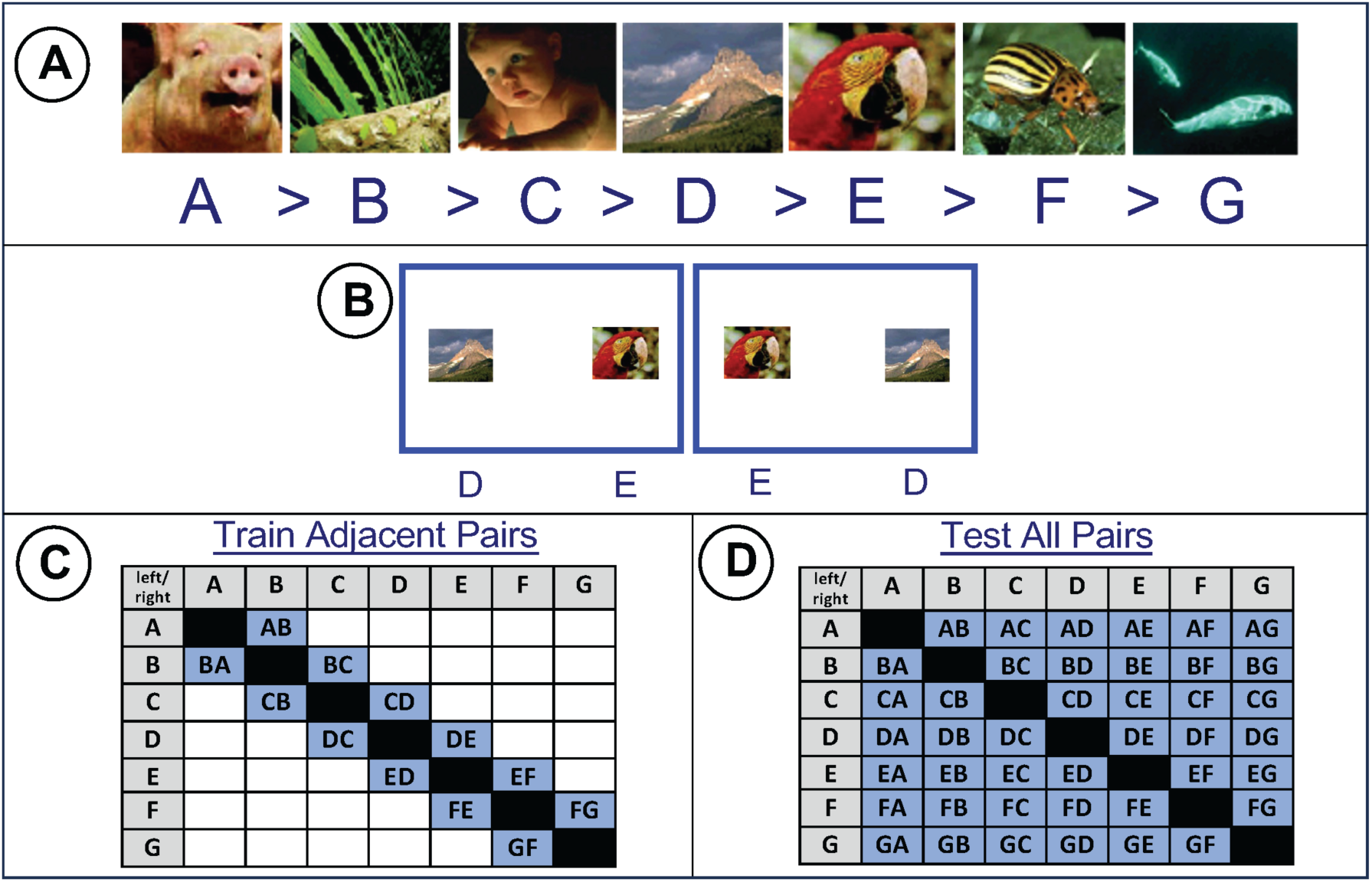
Transitive Inference Transfer Task. This task tests subjects’ ability to learn the implied serial order of a set of images from pairwise presentations. **A.** Set of 7 randomly selected images arranged in an ordered list (A>B>C>D>E>F>G). A novel set of images was used for each behavioral session. **B.** Choice trials with spatial position counterbalanced. **C.** Adjacent pairs used for the training phase of the session **D.** All pairs used for the transfer phase.

On each trial (**Fig. 1B**), the subject first fixated a small target in the center of the screen. Following a random delay (0.4 sec to 1.2 sec, positively skewed with a mean of 0.5 sec), the fixation point was extinguished, and two stimuli appeared equidistant from the center of the screen in opposite directions. To receive a reward, subjects had to select the stimulus with the lower rank by making a saccadic eye movement to the selected stimulus within 1.5 sec of stimulus onset and fixating for a random interval (0.4 to 0.6 sec, uniformly distributed). When this interval elapsed, auditory feedback was delivered indicating that the response was either correct (high tone, 880 Hz) and would be rewarded, or was incorrect (low tone, 440 Hz). To receive the reward on correct trials, subjects had to maintain fixation for another interval (0.35 to 0.65 sec, uniformly distributed), after which the screen went dark and fluid rewards (juice or water drops) were delivered through a sipper tube.

Each session was divided into three phases: single target presentations, training, and testing. During the single target phase, only one stimulus, selected at random, was shown during each trial. The stimulus was shown on the left or right side of the screen and the NHP responded by making a saccadic eye movement to fixate the stimulus. There were approximately 5 blocks of single target presentations, or about 70 trials with left/right counterbalancing. Single target trials provided estimates of simple reaction times.

In the training phase of the task, adjacent pairs of stimuli were presented. Each pair comprised two stimuli whose ordinal positions were adjacent in the list (A/B, B/C, C/D, etc, **Fig. 1C**). To receive a reward, the subject was required to make a saccadic eye movement to the stimulus with the lower rank (e.g. A from A/B). For a 7-item list, there are 6 adjacent pairs. With positional counterbalancing, this resulted in training blocks that were 12 trials in length. This phase generally lasted 20 blocks (i.e. 240 trials in total).

In the testing phase of the task, all possible combinations of the 7 items were presented (**Fig. 1D**). The subjects were still required to saccade to the lower ranking stimulus. This phase tested the subjects’ ability to transfer knowledge about the list order gained during the adjacent pairs phase to novel, non-adjacent pairs. For a 7-item list, there are 21 possible pairs (6 adjacent, 15 non-adjacent). With counterbalancing of the stimulus positions, this resulted in blocks of 42 trials. This phase generally lasted for 5-10 blocks (240-420 trials).

Each pair of stimuli could be identified in terms of the ranks of the individual stimuli they included. Pairs could be also described in terms of “symbolic distance,” a metric that is calculated by taking the difference between the individual stimulus ranks (D’Amato & Colombo 1990). For example, since the pair B/C consists of items that are adjacent in the list (i.e. ranks 2 and 3, respectively), the symbolic distance is 1, while the pair A/G (ranks 1 and 7) has a symbolic distance of 6.

Terminal pairs are those that contain one or both of the terminal items (A,G). Because these items have different outcome associations than non-terminal items (A is always correct, G is never correct, all other items are correct on half of the trials), they can skew the observed results and are excluded from some analyses.

### Sequential Probability Ratio Test (SPRT)

The Sequential Probability Ratio Test (Wald 1945, Wald & Wolfowitz 1948) forms the theoretical basis of Drift Diffusion Models (DDMs). We used the SPRT to simulate evidence accumulation using parameters estimated directly from the behavioral data. This involved two assumptions: First that the subject’s relative preference for each item is represented internally as a likelihood distribution on a one-dimensional scale and, second, that the mean and variance of each item’s likelihood can be modeled as a binomial probability distribution whose parameters are related to choice probability.

To estimate the parameters of the binomial likelihood distribution for each item, every choice of that item was scored as a success and every time it appeared but was unchosen was counted as a failure. The binomial distribution for each item thus represented the likelihood of choosing that item when it was presented. The set of binomial likelihood functions for all 7 items represents their relative position along a “mental line” (Riley 1976, Prado et al., 2012, Bonnefond et al., 2014, Jensen et al., 2015, Ramarat et al., 2023, Gazes et al., 2023, Di Antonio et al., 2024). The item positions reflect relative preference calculated from both correct and incorrect trials.

The likelihood functions allowed us to simulate evidence accumulation for each pair and its dependence on symbolic distance. For any pair of items, e.g. E and F, a random deviate, x, was drawn from the probability density, P(x|E,EF), for the correct item (E) in the pair (EF), where P(x|E,EF) = L(E|x,EF). Then the log of the likelihood ratio was computed, e.g. log[L(E|x,EF) / L(F|x,EF)]. The log of the likelihood ratio (LLR) for each sample corresponds to the momentary evidence in drift diffusion models. The probability density was repeatedly sampled and the LLRs were summed until they reached one of two thresholds (boundaries) that represented the amount of evidence accumulated prior to correct and incorrect choices. The repeated sampling and summing of momentary evidence simulates a drift-diffusion process. The number of samples needed to reach threshold is proportional to reaction time. The proportion of trials in which the correct boundary is reached first is an estimate of performance accuracy (percent correct).

### Generalized Drift Diffusion Modeling (GDDM)

Behavioral data were fit to a Generalized DDM using the PyDDM toolbox (Shinn et al., 2020). We used several different approaches to understand the effect of each variable on the dynamics of the GDDM model in relation to serial learning. In all approaches, the data to be fit were reaction time and accuracy segregated by symbolic distance or trial block. The two conditions (symbolic distance or trial block) were fit separately. Parameters were estimated for each NHP subject individually.

#### Standard model

In the first, “standard,” approach, a minimal version of the GDDM was implemented with four parameters: drift rate (rate of evidence accumulation), boundary level (amount of accumulated evidence required for a decision), noise (instantaneous variability in drift rate), and non-decision time. The time evolution of the decision variable, *x* (accumulated evidence), was given by:

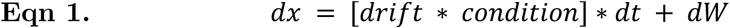

“*Drift*” is the drift rate parameter. “*Condition*” is either symbolic distance or trial block, depending on how the data are segregated, and is coded as an integer; [1…,6] for symbolic distance or [1,…,8] for trial block. “*W*” is the momentary noise represented as a Weiner process with standard deviation = 1. We fixed the noise parameters to *mu* = 0 and *sigma* = 1.0 to avoid redundancy with the boundary height parameter as scaling noise variability is equivalent to scaling the boundary height. The time resolution (dt) was 10 msec and the overall duration was 1.0 sec.

For this approach, the fitting algorithm searched for a single set of parameters (drift rate, bound, and non-decision time) that fit the data within each condition. All parameters except for noise were allowed to vary freely within a wide range (**Table 1**). The drift rate was the product of the scalar drift parameter and the condition index, and therefore varied linearly with condition. The boundary was constant both in time, and across conditions. The same parameter ranges were used for all NHP and all conditions.

**Table 1.** PyDDM parameter ranges.

| Table 1. PyDDM parameter ranges. |  |  |  |  |  |  |
| --- | --- | --- | --- | --- | --- | --- |
| Range | Drift | Boundary | Noise | Leak<br>(if present) | Tau | Non-decision |
| Standard model |  |  |  |  |  |  |
| min | 0.0 | 0.1 | 1.0 | 0 | - | 50 msec |
| max | 10.0 | 10.0 | 1.0 | 100 | - | 400 msec |
| Collapsing bound model |  |  |  |  |  |  |
| min | 0.0 | 0.3 | 1.0 | -50.0 | 0.01 | 50 msec |
| max | 5.0 | 3.0 | 1.0 | 50.0 | 10 | 400 msec |

#### Leaky integration

In the standard approach, there was no loss of evidence over time. However, evidence integration can be imperfect. Thus, in a second, “leaky integration” approach, imperfect integration was modeled by adding a leak term to the standard model:

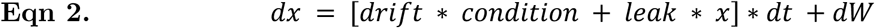

The *leak* is a constant that multiplies accumulated evidence (*x*) at each moment. If the leak term is negative, the evidence tends to decay exponentially toward zero. If the leak is positive, it acts like a gain factor that causes the evidence to increase exponentially (unstable integration).

#### Collapsing bound

In both the standard and leaky integration approaches, the decision boundary remained constant while evidence was being accumulated. To understand the effect of allowing the boundary to change over time, a “collapsing bound” approach was tested. In this approach, the bound collapsed exponentially, thus introducing a sixth parameter, *tau*, the collapse rate. In this approach, the equation for momentary evidence was the same as **Eqn 1**, but the time-dependence of the boundary was given by:

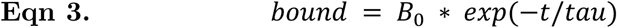

Where *B_0_* is the initial boundary level at *t* = 0. This model was tested both without a leak parameter (using **Eqn 1** for evidence accumulation) and with a leak (using **Eqn 2**).

### Simulations

To illustrate how each model parameter affected dynamics and performance, simulations of the standard, leaky integration, and collapsing bound models are shown in **Fig 2**. The left panel illustrates the time evolution of accumulated evidence for a range of drift rates (signal strengths). The center panel shows the reaction time histograms for correct and error trials. The right panel shows accuracy and reaction time. Non-decision time was set to zero in all simulations.

**Figure 2.**
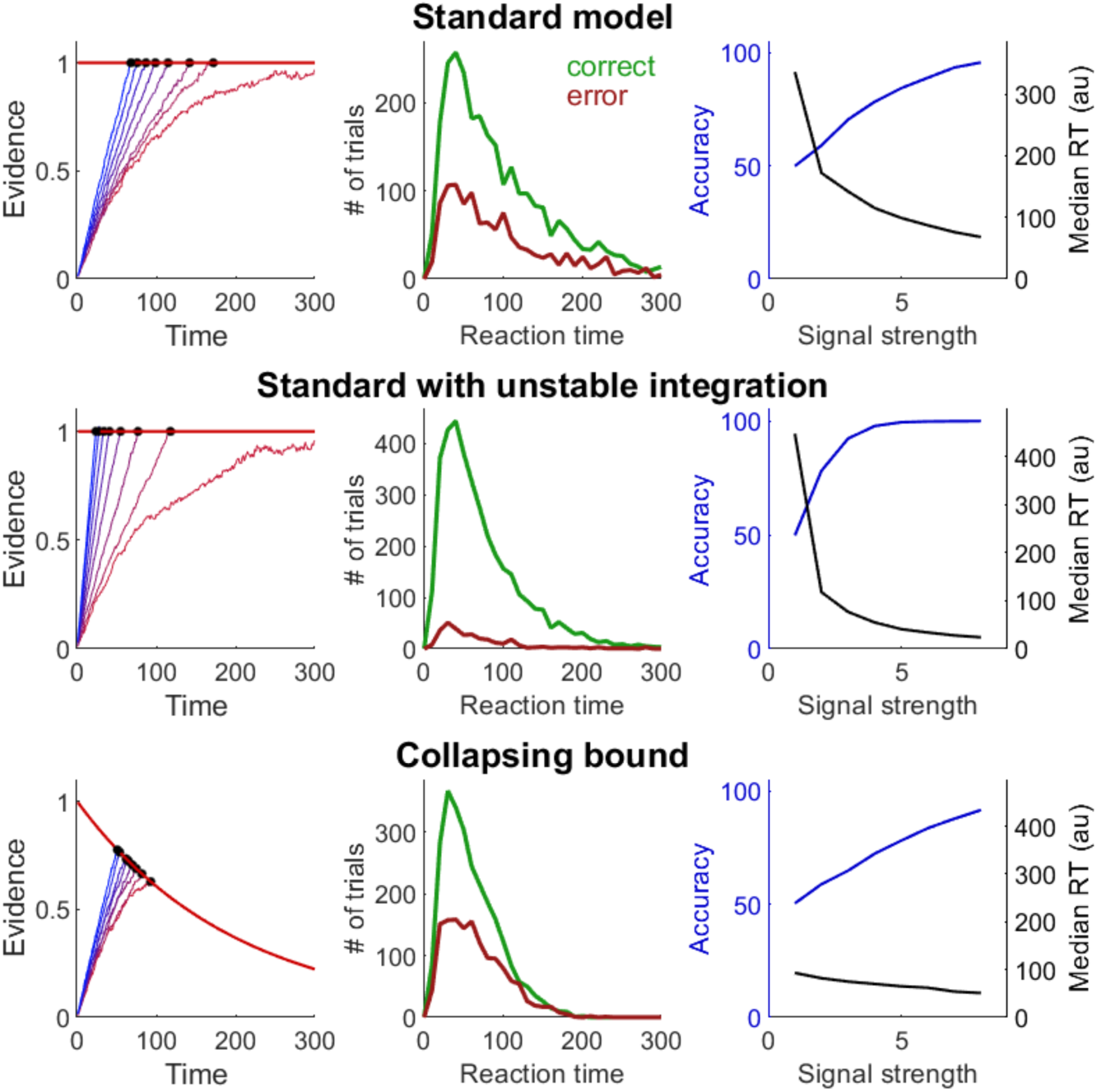
DDM Simulations. **Top:** Standard model with constant bound and no leak. Left: red traces represent weaker signal strengths, blue is stronger signal. Center: reaction time histograms for correct (green) and error (red) trials for a single, moderate signal strength. Right: Accuracy (percent correct) and reaction time as a function of signal strength. **Middle:** Standard model with leak. **Bottom:** Collapsing bound model without leak.

For the standard model (**Fig 2**, top row), low signal strength was associated with chance performance accuracy and long reaction times. The addition of a positive leak term (**Fig 2**, middle row) restricted the range of reaction times by speeding up evidence accumulation, but also reduced accuracy for moderate signal strengths. The collapsing bound model without a leak term (**Fig 2**, bottom row) performed similarly to the leaky integration model but was more effective in reducing long reaction times and produced more symmetric reaction time distributions.

## Results

### Behavioral Performance with Eye Movement Responses

Performance was quantified in terms of accuracy (percent correct) and response time (time from stimulus onset until choice saccade onset). A total of 203 behavioral sessions were recorded (F: 85, H: 98, L: 20 sessions). Each session started with a novel set of pictures that were arranged into a list by assigning a rank order to each picture (**Fig. 1A**). The ranking was initially unknown to the subject but was learned over the course of the session.

Reaction times (RT) as a function of trial number for all sessions completed by NHP H are shown in **Fig. 3A**. The three phases of the task are color-coded (single target trials in gray, adjacent pairs in blue, all pairs in magenta). There was an abrupt shift in the RT distribution (**Fig. 3B**) at the transition from single stimuli (median RT 192 msec) to stimulus pairs (median RT 270 to 275 msec).

**Figure 3.**
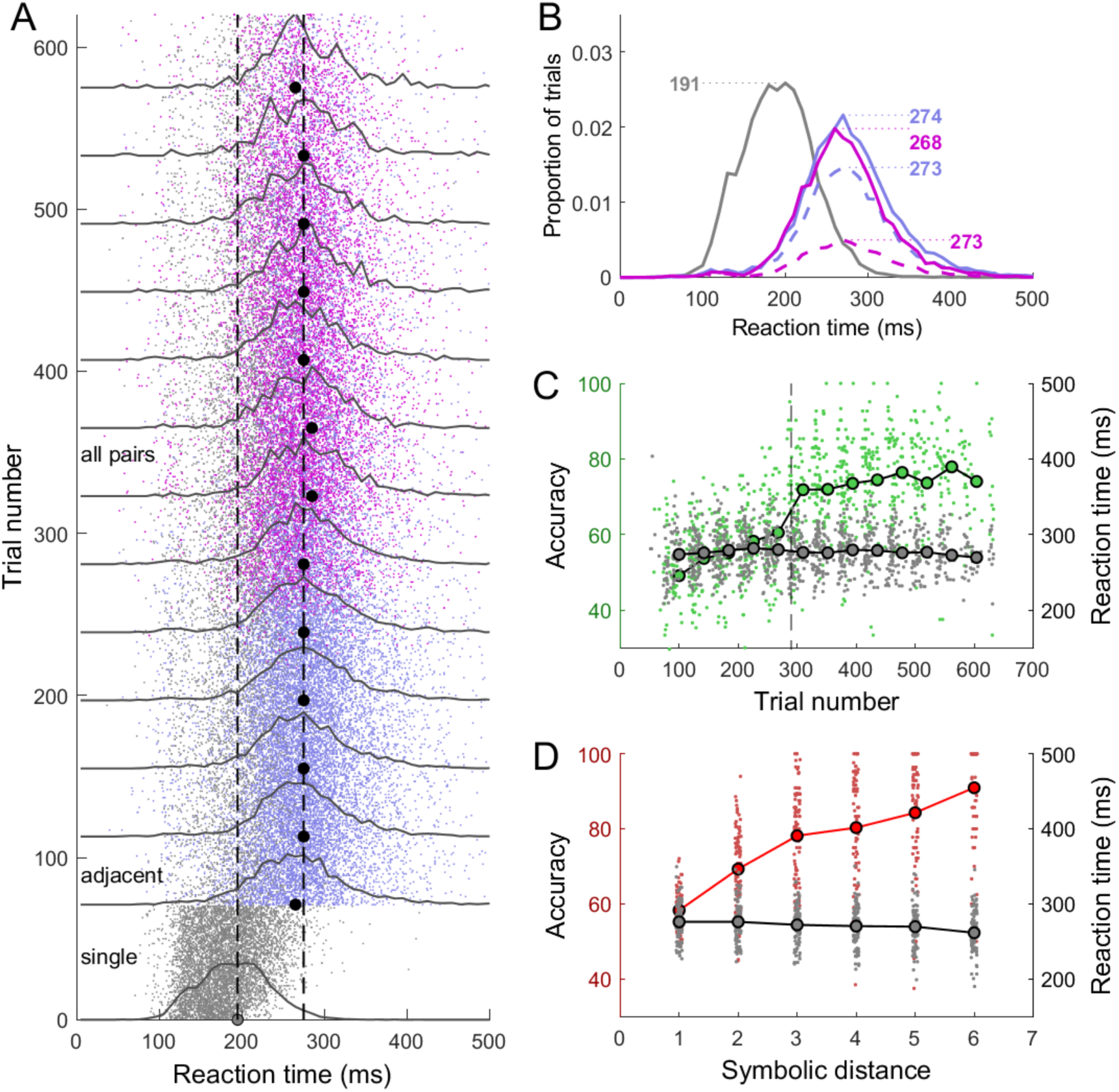
Behavioral Performance on TI transfer task. **A.** Reaction time vs trial number for NHP H. The colors represent the three task phases (grey – presentation of single items, cyan – adjacent pairs, red – all pairs). The gray dot on the x-axis and the thin dashed line is the median RT for single target trials. The black dots are the median RTs for choice trials binned in 50 trial segments. The thick dashed line is the median over all choice trials. **B.** Reaction time histograms. Colors as in A. For choice trials, solid lines are correct responses and dashed lines are incorrect. **C.** Accuracy (green) and reaction time (gray/black) during learning. Dashed vertical line is transition from training to testing phase. **D.** Accuracy (red) and reaction time (grey/black) as a function of symbolic distance.

Adjacent-pair (training phase) and all-pair (testing phase) trials were grouped into blocks of 42 consecutive trials to obtain robust estimates of RT (**Fig. 3A**). The shape and median value of the RT distributions remained constant across trial blocks. The RT distributions were similar in shape for correct (**Fig 3B** solid lines) and incorrect (dashed lines) trials, and for adjacent (**Fig. 3B**, blue lines) and non-adjacent pairs (magenta lines).

The effect of learning on accuracy and RT for this NHP is shown in **Fig. 3C**. During adjacent pair training, logistic regression showed a significant effect of trial number on accuracy for all NHPs (F: *p* = 0.03, df = 15,206; H: *p* < 0.0001, df = 16,012; L: *p* < 0.0001, df = 6658). Logistic regression coefficients for accuracy versus trial number were similar for all NHP (0.096 for NHP F, 0.07 for NHP H, 0.11 for NHP L). The effect of trial number on RT, as assessed by linear regression, was also significant (*p* < 0.0001 for all NHPs). However, the slopes of the regression lines were shallow, amounting to an increase of 5 to 7 msec for every 100 trials.

During all-pairs testing, the relationship between trial number and accuracy remained significant (*p* < 0.0001 for all NHP) and the RT trend turned slightly negative (−0.5 msec per 100 trials) and was significant for 2 NHP (*p* < 0.002). However, the relationship was inconsistent - negative for one NHP (H: decrease of 1.6 ms/400 trials), and positive for the other (L: increase of 4 ms/400 trials).

Symbolic distance (SD, the difference in rank between the two stimuli in each pair) had a strong effect on accuracy (**Fig. 3D**). Logistic regression coefficients for SD were consistent across NHP (0.35 for NHP F, 0.28 NHP H, 0.22 for NHP L). For RT, there was a small but significant reduction with SD for both NHPs (F: slope = −1.5 msec, *p* < 0.0001; H: −2.9 msec, *p* < 0.0001; L: −0.33 msec, *p* = 0.02). The results overall indicate a strong effect of trial number and SD on performance accuracy, but little effect on response time.

#### Accuracy and Reaction Time at Transfer

The transition from adjacent pair training to all-pairs testing was not signaled to the NHP while they were performing the task. Rather, non-adjacent pairs were simply added into the mix of trials. The start of the testing phase is defined as the occurrence of the first non-adjacent pair after adjacent-only training. Performance accuracy showed an abrupt effect of symbolic distance from the first testing trial onward. **Fig. 4A** shows training and testing accuracy averaged over all sessions for the 3 NHP. After ~200 training trials, accuracy was close to 60% correct. At the start of all-pairs testing, accuracy for adjacent pairs (**Fig. 4A** black lines follow the same trajectory as training. However, at the beginning of testing, accuracy for non-adjacent pairs (**Fig. 4A** colored lines) was superior to adjacent pairs for each symbolic distance even though all non-adjacent pairings were novel. We have documented this remarkable jump in performance at transfer with other NHP and other testing systems (Jensen et al., 2015). The same pattern of results was obtained even when terminal pairs (those containing items A or G) were excluded (**Fig. 4B**). By contrast, reaction times showed minimal variation across training and testing (**Fig. 4C,D**) To examine performance at transfer more closely, the accuracy (**Fig 4E**) and reaction time (**Fig. 4F**) for each item pair during the first block (42 trials) of testing are shown. During a block of 42 trials, each pair was shown twice; once in each spatial configuration.

**Figure 4.**
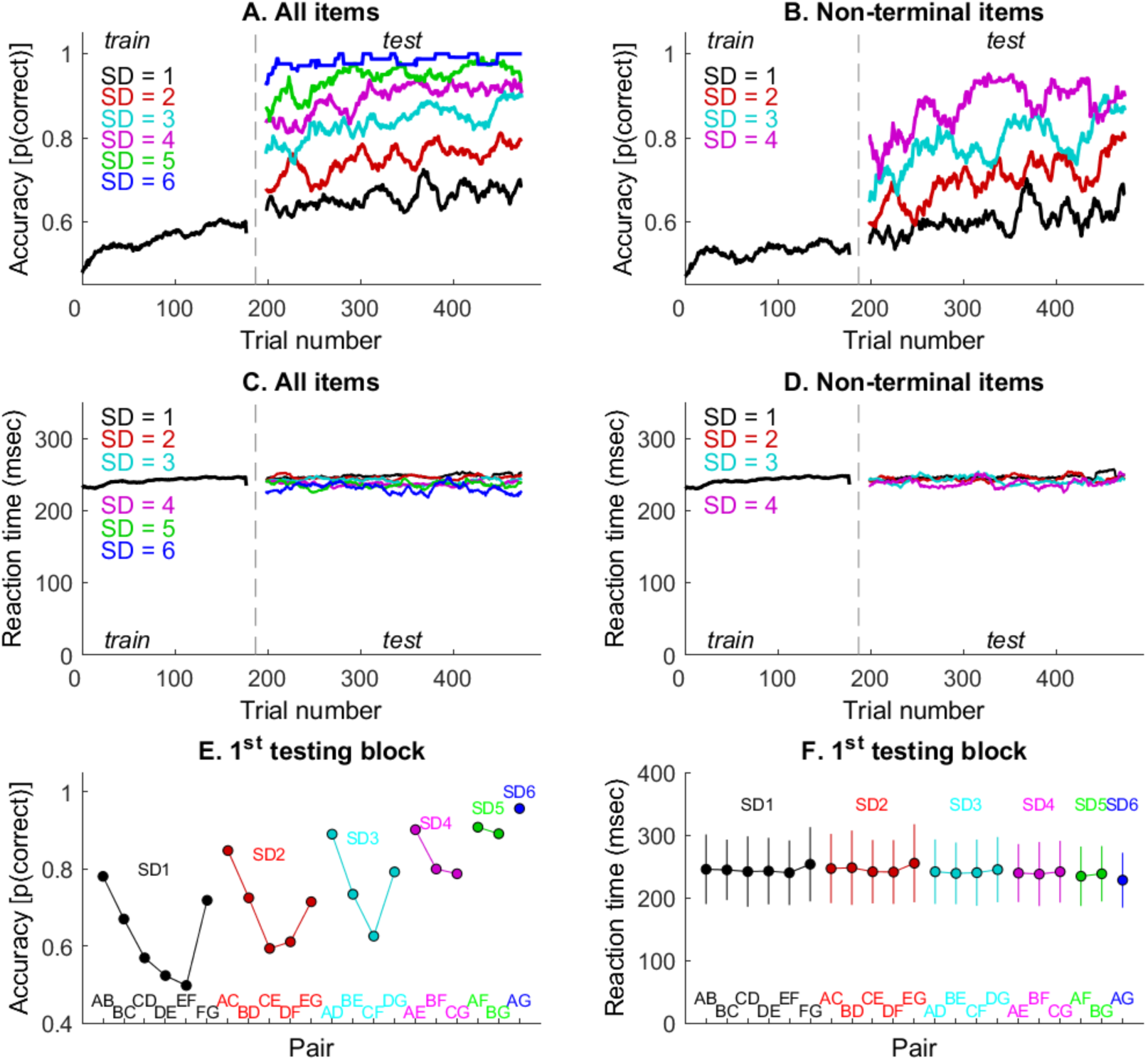
Accuracy and reaction time at transfer. **A.** Accuracy vs. trial during training and testing averaged over all NHP and sessions. **B.** Accuracy for pairs with non-terminal items. **C.** Reaction time during training and test. **D.** Reaction time for critical pairs. **E.** Accuracy for each pair during the first block of testing. **F.** Reaction time for each pair during the first block of testing.

The six test pairs, BD, CE, DF, BE, CF, and BF are “critical” in the sense that they did not occur during training nor do they contain terminal items. Above chance performance on these pairs, as well as the overall effect of symbolic distance, is considered evidence that NHP relied on transitive relations in making their choices. For example, during training, B and F were chosen almost equally often from the pairs BC and FG (in fact F was correctly chosen from FG, and therefore rewarded, more often than B was correctly chosen from BC). Yet, during the first block of testing, B was chosen over F in roughly 80% of the BF trials.

#### Speed-accuracy trade-off

Although there was not a clear relationship between accuracy and choice reaction time when data were combined over sessions (**Figs 3C,D**), a more fine-grained analysis revealed subtler trends. In this analysis, we calculated the residual accuracy and average reaction time in each block of 42 consecutive trials during the adjacent-pair training and all-pairs testing. “Residual” is here defined as the accuracy or average reaction time within each block of 42 trials normalized by subtracting the averages of each of those measures calculated over all choice trials for the entire session. By compensating for session-to-session variability, the residuals provided estimates of the change in accuracy per unit change in reaction time that could be combined across sessions.

**Figs. 5A,B,C** show the improvement in accuracy with increasing reaction times during the training phase of the task (first 200 choice trials of each session) for all three NHP. Each data point represents a single 42-trial block from one session. For every additional millisecond of reaction time, there was an average increase in performance ranging from 0.24 to 0.43 percent correct. The correlation between accuracy and RT was significant for each NHP. This is opposite to the typical pattern for perceptual decisions where increasing accuracy is associated with shorter reaction times. During the subsequent testing phase (the first 200 trials after end of training)(**Fig. 5 D,E,F**), no significant relationship between accuracy and reaction time was found for any NHP.

**Figure 5.**
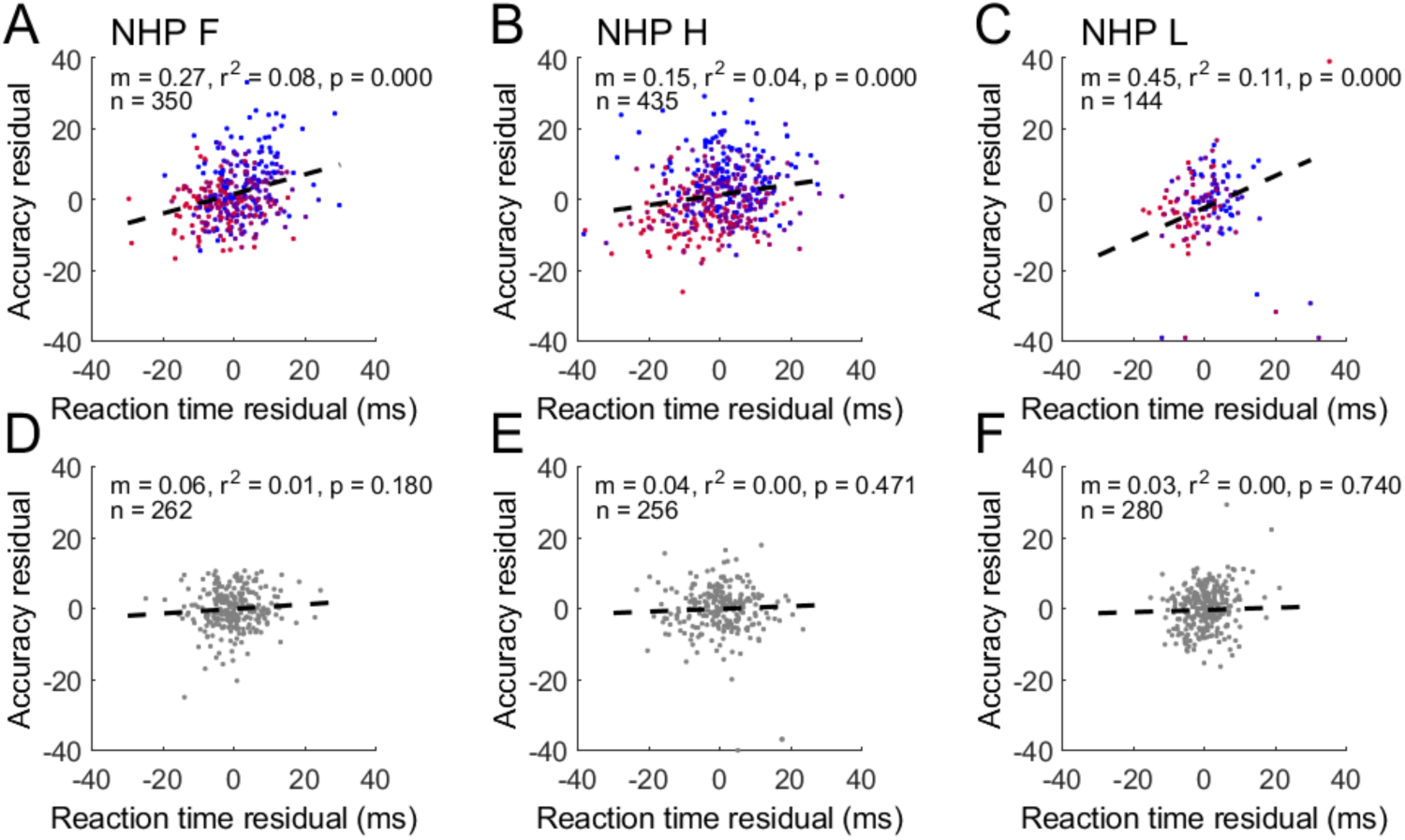
Speed-accuracy trade-off. **A,B,C.** Improvement in accuracy v. increase in RT during training phase for each NHP. ‘m’ is the linear regression slope, ‘r2’ is the coefficient of determination, ‘p’ is the significance of the correlation (Pearson), and ‘n’ is the number of 50-trial blocks per session times the number of sessions. The color gradient indicates trial blocks that were earlier in training (red) vs. later (blue). **D,E,F.** All-pairs testing phase.

During the all-pairs testing phase, we sorted trials into six groups by symbolic distance (SD), rather than by trial block. When this was done, a significant negative relationship was found between the change in accuracy and increasing reaction time (**Fig. 6 A,B,C**); trials with larger SD had higher accuracy and shorter RTs.

**Figure 6.**
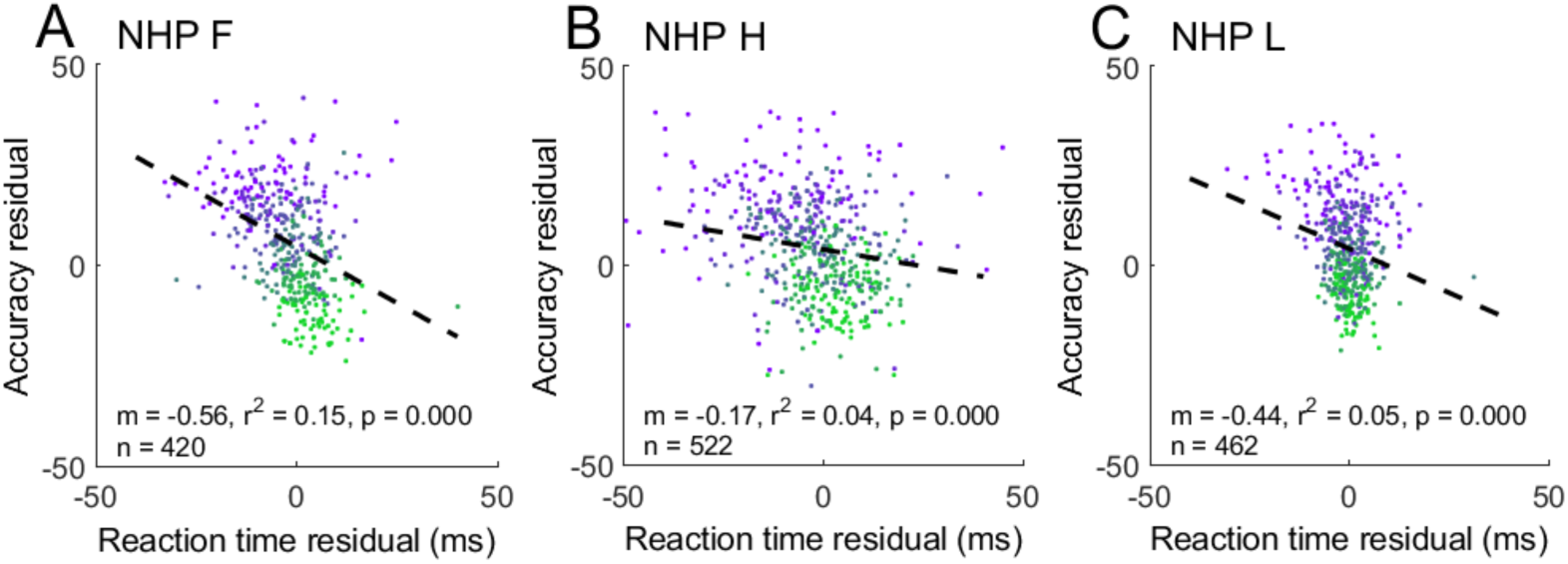
Speed-accuracy trade-off for trials grouped by symbolic distance. **A,B,C.** Decrement in accuracy v. increase in RT during testing phase for each NHP. ‘m’ is the linear regression slope, ‘r2’ is the coefficient of determination, ‘*p*’ is the significance of the correlation (Pearson), and ‘n’ is the number of symbolic distances times the number of sessions. The color gradient indicates smaller (green) vs. larger (magenta) symbolic distances.

### Behavioral Performance with Reaching Movements

Three additional NHP (O, Q, R) were trained to perform the TI task by reaching to touch one of the two images presented on a touchpanel. The task was otherwise identical to the version with eye movement responses. NHP O completed 25 sessions with an average of 540 trials per session, Q completed 112 sessions with an average of 459 trials, and R completed 42 sessions with an average of 441 trials. **Fig. 7A** shows performance averaged across all sessions and NHPs for the last 100 trials before and 200 trials after transfer. **Fig. 7B** shows touch reactions times. The results were nearly identical to those obtained with eye movement responses (**Fig. 4A,C**) except that reaction times for touch responses were overall roughly 350 msec longer than for eye movements. Just as for eye movements, decision accuracy showed a sudden improvement and symbolic distance effect at transfer while reaction times remained flat with a very small symbolic distance effect. Faster responses tended to be more accurate both within and across symbolic distance (**Fig. 7C**), though the association was weak.

**Figure 7.**
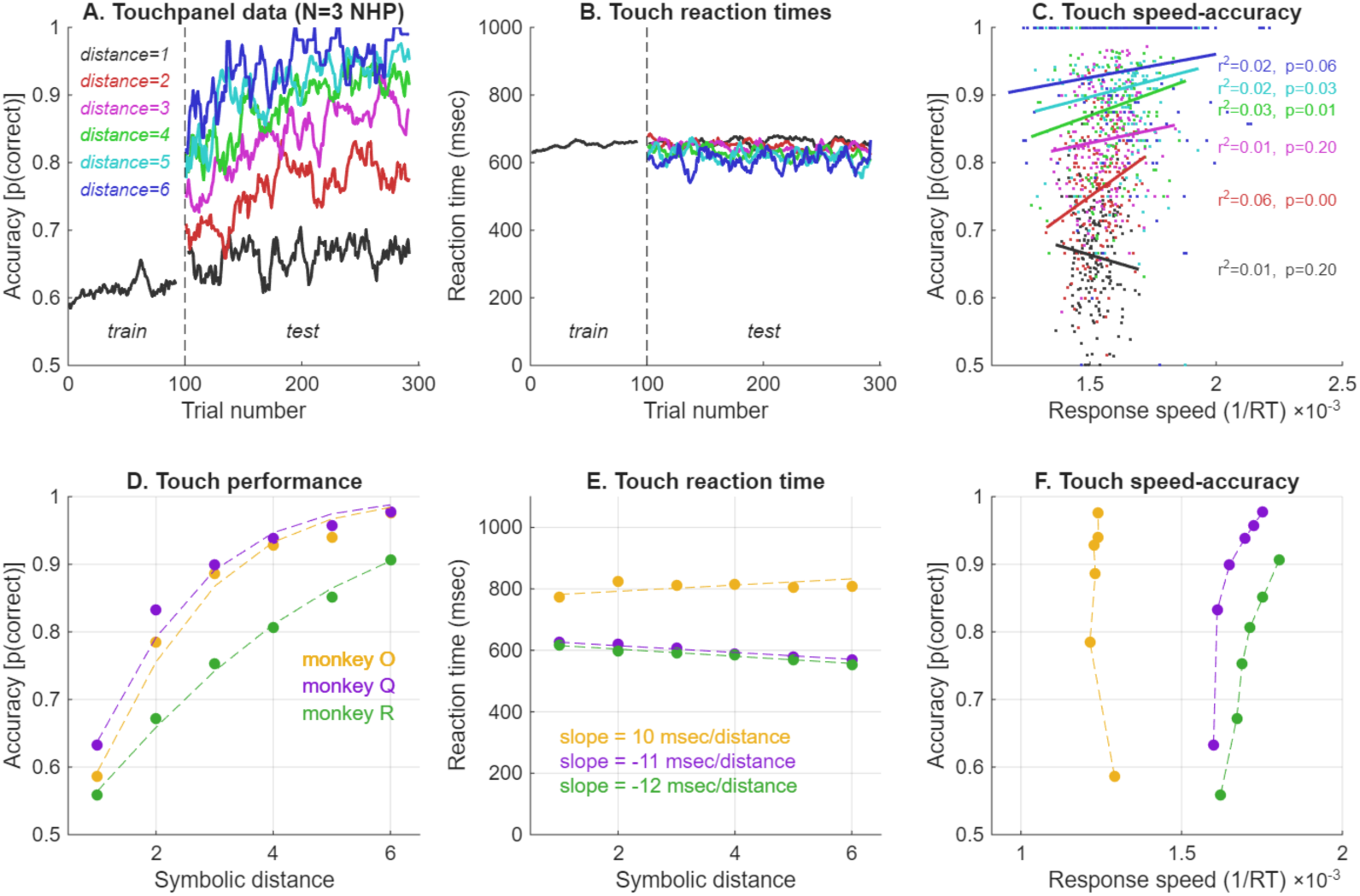
Touchpanel performance. **A.** Decision accuracy averaged over NHP(n=3) and sessions (n=179). **B.** Reaction time averaged over NHP and sessions. **C.** Speed-accuracy trade-off. **D.** Accuracy for each NHP, dashed lines are logistic regression fits. **E.** Reaction time for each NHP, dashed lines are linear regression fits. **F.** Speed-accuracy trade-off for each NHP.

The results for individual NHP confirmed a strong effect of symbolic distance on decision accuracy (**Fig. 7D**), but little effect on reaction time (**Fig. 7E**). One NHP showed an increase in RT with symbolic distance while the other two showed decreases, but the slopes in all 3 cases were quite shallow, averaging 2.1%. For 2 NHP, the most accurate decisions were about 10% faster than the least accurate, while the third animal showed a slight reduction of response speed with increasing accuracy (**Fig. 7F**)

### Rate of evidence accumulation estimated from sequential sampling

An optimal method for making binary decisions is to repeatedly sample the relative likelihood of the two alternatives (Wald & Wolfowitz 1948) until the sum of the log-likelihood ratios reaches a predetermined value (bound). We applied this method to determine the optimal rate of evidence accumulation (i.e. diffusion) in the TI task. Because the NHPs made a binary choice on each trial, it is reasonable to model their choices using a set of binomial likelihood distributions, one for each item (Jensen et al., 2015). The parameters of those distributions (success rate, *p,* and number of trials, *n*) correspond to the number of times each item was chosen relative to th e number of times it was presented (choice probability). This is demonstrated for the pair EF in **Fig. 8A** which shows the likelihood functions estimated from the pattern of choices made at the end of the training phase of the task averaged over all 3 NHP.

**Figure 8.**
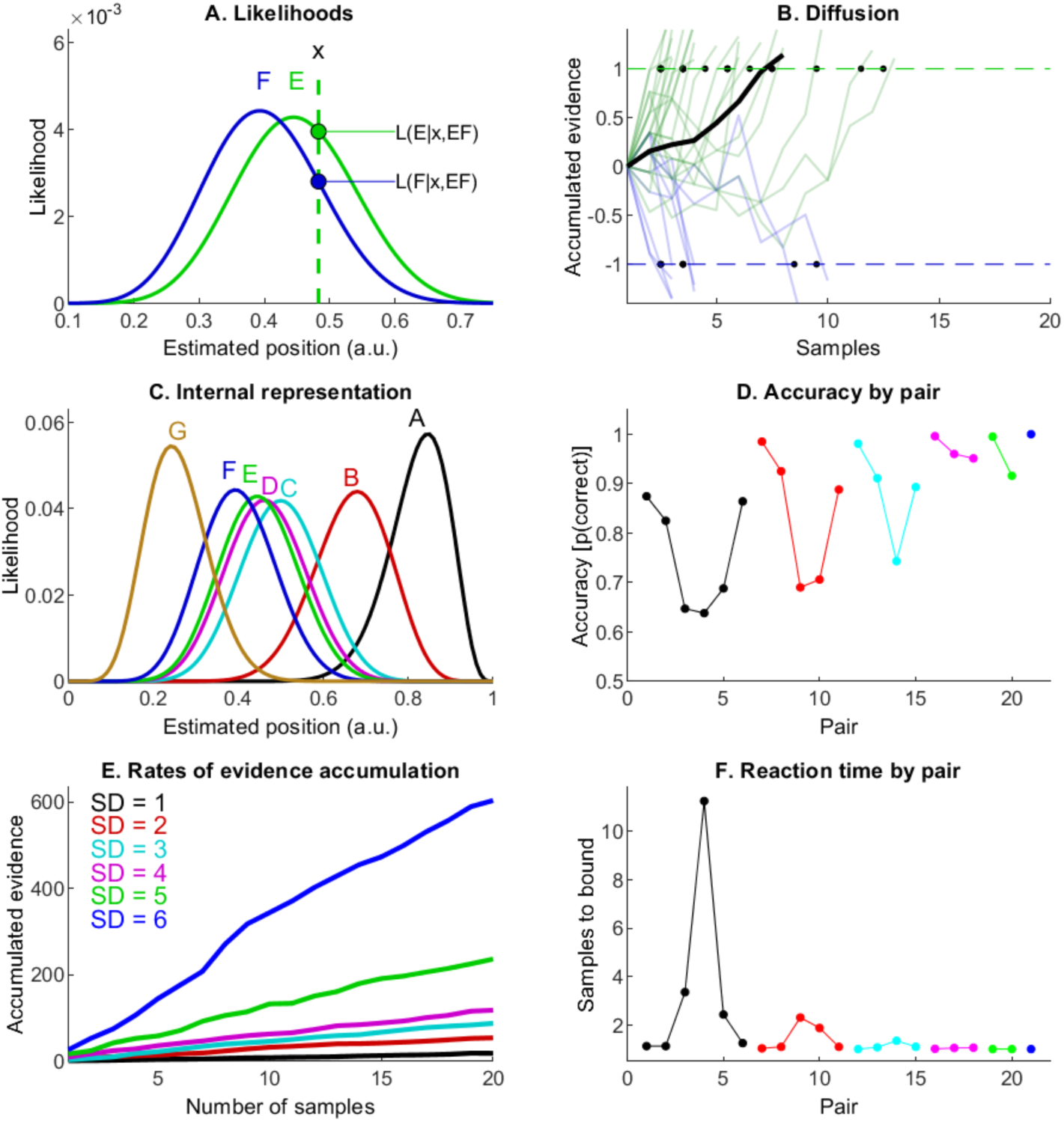
Sequential probability ratio test (SPRT). **A.** Binomial likelihood functions for items E and F. **B.** Accumulation of evidence for the pair EF based on repeated sampling of relative likelihood. Upper bound (green dashed line) corresponds choices of E, lower bound (blue) corresponds to choosing F. Solid black line is the average accumulated evidence. **C.** Binomial likelihood functions for all items. **D.** Accuracy for each pair predicted by SPRT. **E.** Evidence accumulation for different symbolic distances. **F.** Reaction time (samples to bound) predicted by SPRT.

The likelihood functions for each item were used to simulate a sequential sampling decision process (**Fig. 8B**) with constant bounds. The rate of evidence accumulation and the momentary noise were determined by the means and widths of the likelihood functions for E and F, combined with random draws from the likelihood function for E. The upper and lower bounds were set to approximate the observed probability of choosing E from the pair EF.

The likelihoods for all 7 items, estimated from choice probabilities at the end of testing, are shown in **Fig. 8C**. We propose that this set of functions forms an internal representation of the ordering of the TI list and the degree of uncertainty associated with the relative position of each item (Jensen et al., 2015). Simulations were run to estimate the rate of evidence accumulation for each pair during the testing phase. Using a fixed bound, we could estimate the accuracy and reaction time (number of samples to bound) for each pair. The pattern of accuracy vs. pair (**Fig. 8D**) was qualitatively similar to the behavioral data (**Fig. 4C**), thus validating the approach, at least in part.

Evidence accumulation rates were combined over pairs to estimate the rate of evidence accumulation as a function of symbolic distance (**Fig. 8D**). Evidence accumulation was approximately linear with the number of samples, reflecting the fact that, on average, each sample provided the same amount of momentary evidence regardless of when it was drawn. The rate of accumulation (drift rate) increased with symbolic distance.

It is obvious that there is no constant boundary level that could be reached after a fixed number of samples for all symbolic distances. This is confirmed by plotting the number of samples (a proxy for reaction time) needed to reach a bound that was fixed for all pairs (**Fig. 8F**). Reaction times produced by this version of the SPRT were inversely related to accuracy, a result that does not match the behavioral data (**Fig. 4D**).

In general, the accuracy and reaction times obtained with a simple but theoretically well-motivated sequential sampling model are at odds with the experimental data and suggest that a fuller treatment is needed. We therefore turned to the PyDDM package to test more elaborate models.

### Generalized DDM Modeling of Decision Accuracy and Reaction Time

For eye movements, choice reaction times during the TI transfer task were generally under 300 msec even though subjects were allowed up to 1500 msec after stimulus onset to respond by making a saccadic eye movement to the target or distractor. Choice RTs averaged only 35 msec longer than simple RTs for NHP F (215 vs. 180 msec median RT), 85 msec for NHP H (275 vs. 190 msec), and 30 msec for NHP L (211 vs. 181 msec). For reaches, RTs were over twice as long as eye movement RTs, raising the question of whether decision dynamics during reaches were stretched in time or merely delayed.

For both eye movements and reaches, RTs remained nearly constant despite learning-related improvements in decision accuracy. RTs were also constant across symbolic distance despite large differences in accuracy. Drift diffusion models (DDMs) with constant bounds and variable drift rate typically show increases in accuracy that are accompanied by shorter reaction times. As the current results did not show this pattern, it raises the question of whether DDMs can fit the behavior exhibited by NHP during the transitive inference task and thus shed light on how decisions are made during TI learning.

To test this, we first fit a generalized DDM with four parameters (drift rate, noise, boundary, and non-decision time, see Methods) to the current behavioral data using the PyDDM package (Shinn et al. 2020). An example of the PyDDM fits for data segregated by outcome (green = correct, red = error) for NHP L is shown in **Fig. 9**. The fits for the standard model are shown as the green and red curves. These fits were relatively poor. This was confirmed by the variance accounted for (VAC, measured by r^2^), which was generally less than 80%. The fitted parameter values are shown in **Table 2**.

**Figure 9.**
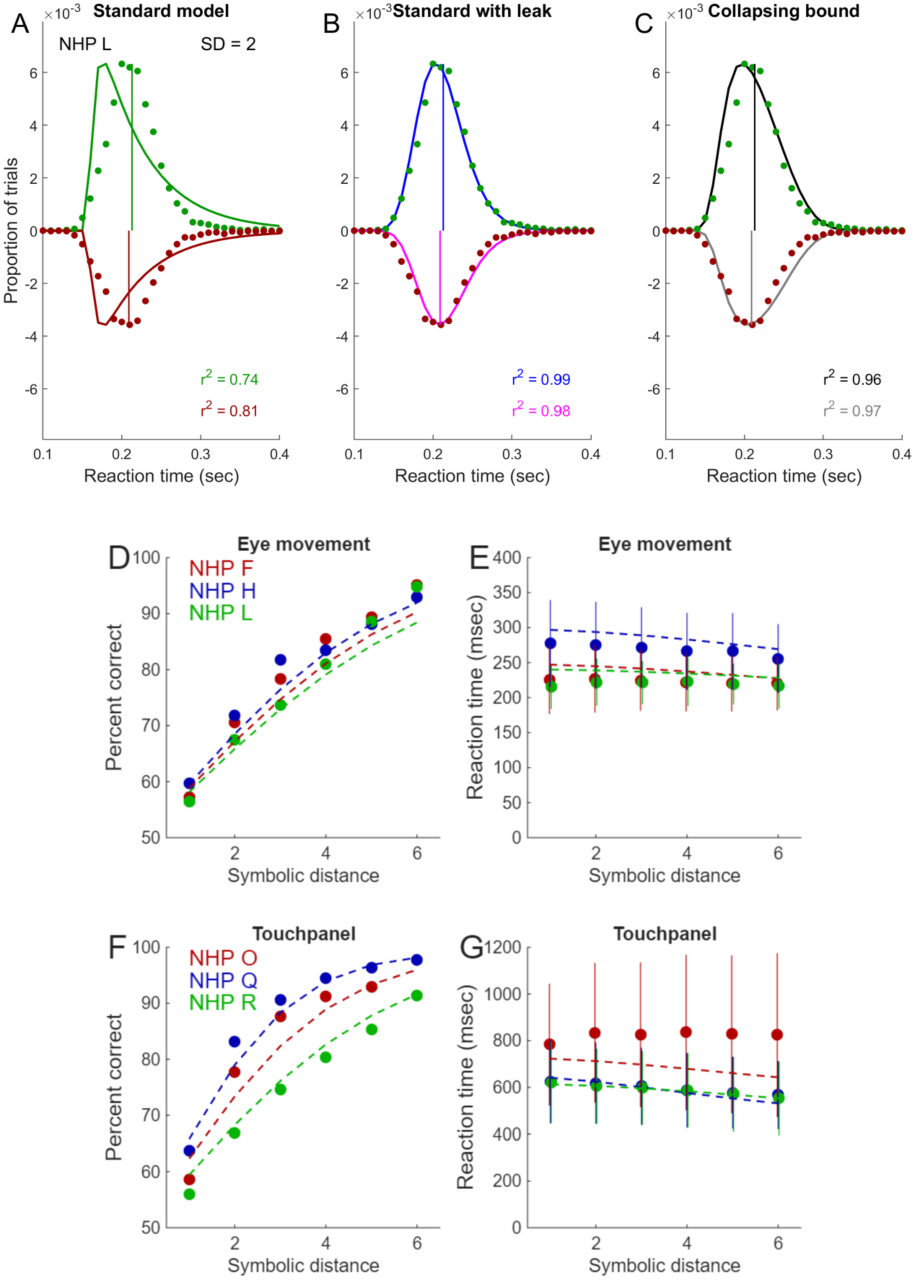
Generalized Drift Diffusion Model fits. Data from all sessions for NHP L with symbolic distance (SD) equal to 2. Green dots are reaction time distributions for correct trials, red dots for error trials. Vertical red and green lines indicate the median RT for each distribution. Green and red curves are fits for correct and error trials, respectively, using the standard model. The correlation between the fits and the data are shown in red and green text. Blue and magenta curves and text are fits obtained with the leaky integration model. Black and grey curves and text are fits obtained with the collapsing bound model. **D-G**. Model 3 (collapsing bound) fits to average accuracy and reaction time for eye movments (**D,E**) and reaching movements (**F,G**) for all six NHP.

**Table 2.** PyDDM fitted parameters for each model, NHP, and condition. For “condition,” SD is symbolic distance. NHP F, H and L responded with eye movements, while O, Q, and R responded by reaching to a touchpanel. Model 1 is the standard model with constant bound and no leak. Model 2 is the standard model with a leak term. Model 3 is the collapsing bound model with no leak. Model 4 is the collapsing bound with leak. BIC is Bayesian information criterion.

| NHP - Condition | Model | Leaky Integration | Collapsing Bound (Tau) | Drift | Bound | Noise | Non-decision | BIC |
| --- | --- | --- | --- | --- | --- | --- | --- | --- |
| NHP F - SD | 1 | - | - | 0.68 | 0.31 | 1.0 | 148 msec | -49407 |
| NHP F - SD | 2 | 29.2 | - | 0.76 | 2.85 | 1.0 | 104 msec | -56392 |
| NHP F - SD | 3 | - | 0.14 | 0.54 | 0.68 | 1.0 | 118 msec | -57131 |
| NHP F - SD | 4 | 26.9 | 7.9 | 0.72 | 2.29 | 1.0 | 102 msec | -56379 |
| NHP H - SD | 1 | - | - | 0.67 | 0.35 | 1.0 | 177 msec | -18819 |
| NHP H - SD | 2 | 20.1 | - | 0.71 | 2.98 | 1.0 | 106 msec | -23881 |
| NHP H - SD | 3 | - | 0.16 | 0.58 | 0.89 | 1.0 | 135 msec | -23923 |
| NHP H - SD | 4 | 20.2 | 6.8 | 0.71 | 2.79 | 1.0 | 110 msec | -23853 |
| NHP L - SD | 1 | - | - | 0.72 | 0.30 | 1.0 | 156 msec | -13061 |
| NHP L - SD | 2 | 42.4 | - | 0.86 | 2.7 | 1.0 | 133 msec | -15369 |
| NHP L - SD | 3 | - | 0.09 | 0.60 | 0.59 | 1.0 | 140 msec | -15241 |
| NHP L - SD | 4 | 43.0 | 7.2 | 0.86 | 2.83 | 1.0 | 131 msec | -15367 |
| NHP O - SD | 1 | - | - | 0.57 | 0.51 | 1.0 | 490 msec | -3182 |
| NHP O - SD | 2 | 7.7 | - | 0.57 | 1.22 | 1.0 | 444 msec | -4105 |
| NHP O - SD | 3 | - | 0.44 | 0.50 | 0.99 | 1.0 | 438 msec | -4658 |
| NHP O - SD | 4 | -1.0 | 0.41 | 0.50 | 0.88 | 1.0 | 448 msec | -4650 |
| NHP Q - SD | 1 | - | - | 0.71 | 0.52 | 1.0 | 405 msec | -16163 |
| NHP Q - SD | 2 | 8.5 | - | 0.71 | 2.14 | 1.0 | 307 msec | -21485 |
| NHP Q - SD | 3 | - | 0.41 | 0.64 | 0.97 | 1.0 | 350 msec | -22057 |
| NHP Q - SD | 4 | 7.8 | 1.53 | 0.68 | 2.41 | 1.0 | 297 msec | -21512 |
| NHP R - SD | 1 | - | - | 0.38 | 0.51 | 1.0 | 360 msec | 342 |
| NHP R - SD | 2 | 8.1 | - | 0.44 | 3.0 | 1.0 | 222 msec | -3045 |
| NHP R - SD | 3 | - | 0.42 | 0.36 | 1.06 | 1.0 | 299 msec | -3316 |
| NHP R - SD | 4 | 8.05 | 9.8 | 0.44 | 2.93 | 1.0 | 227 msec | -3028 |

#### Effect of Drift Rate

The drift term captures whether the rate of evidence accumulation is sensitive to the stimulus conditions, as indexed by symbolic distance. The fitted parameters indicate an increased rate of evidence accumulation (drift rate) as a function of SD. This is expected as drift rate has a strong effect on accuracy. The drift rate estimates were similar across NHP and movement type.

#### Effect of Leak

The leak term modulates the rate of evidence accumulation. It can be thought of as a gain control that dampens or enhances the effect of drift rate and has a more powerful effect as more evidence is accumulated. Adding a leak term to the standard model improved the fits, as shown by the blue and magenta curves (**Fig. 9**). These fits were substantially better than the standard non-leaky model, resulting in VACs in the range of 88% to 97%. The Bayesian Information Criterion (BIC) scores were also improved (**Table 2**).

The leak term can either slow or accelerate the rate of evidence accumulation depending on its sign. For the standard model with a leak, the fitted leak term was positive, indicating unstable integration. The magnitude of the leak term was quite large such that it played a dominant role in the model’s performance, minimizing the effect of drift rate and boundary.

#### Effect of Boundary

The initial boundary level sets the amount of accumulated evidence needed to render a decision, whereas the collapse rate determines if that amount of evidence is stable over time. The collapsing bound model with no leak provided fits that were as good as the standard (constant bound) model with leaky integration, as measured by VAC and BIC. Adding a leak term to the collapsing bound model had a negligible or deleterious effect on the fits (**Table 2**, compare BIC for models 3 and 4). Models that included a leak term tended to also use a higher boundary setting, indicating a positive relationship between these parameters. There was no indication that boundary levels differed between eye and reaching movements.

#### Effect of Non-decision Time

All NHP had non-decision times that were similar across models. However, there were differences among the NHP. Non-decision times for NHP F and L were on average 30-32 msec shorter than NHP H. It is of interest that the simple RTs for NHP F and L had the same average (181 msec), while H had the longest simple RTs at 192 msec suggesting that simple RTs may be related to non-decision time.

The large difference (roughly 340 msec) in overall RT between eye and reaching movements was reflected in the fitted non-decision times. For reaches, non-decision time averaged 357 (+/- 89 sd) msec across NHP and models, vs. 130 (+/-23) msec for eye movements (t-test t=8.6, p<0.0001, df=22.)

#### Speed-accuracy trade-off

The modeling approaches considered so far have been evaluated on the basis of their ability to fit average accuracy and reaction time data. We have not yet considered the relationship between residual accuracy and reaction time (i.e. the speed-accuracy trade-off) shown in **Figs. 4** and **5**. The striking feature of these trade-offs is that they showed a positive slope as a function of trial block during training and negative slope as a function of symbolic distance during testing.

With a fixed bound, the DDM tends to produce a negative relationship between accuracy and reaction time as indicated by the simulations in **Fig. 2**. One way to turn this into a positive relationship is to allow the boundary to vary. If the boundary height increases with learning, then accuracy and reaction time can increase together as in **Fig. 5A-C**. The relationship between boundary position and signal strength that leads to increasing reaction times can be calculated analytically (Bogacz et al 2006, 2008). These calculations reveal that accuracy is related to the product of drift rate and boundary position, while reaction time is more closely related to their ratio.

### GDDM Modeling of Learning and Transfer

To estimate the effect of within-session learning on model fits, the data within each session were grouped into 8 blocks of 50 consecutive trials spanning the last 200 trials of adjacent-pair training and the first 200 trials of all-pairs testing. Blocks were then combined across sessions. During each session, accuracy improved from chance levels to roughly 80% correct. However, there was a sharp increase in accuracy between the end of training and the onset of testing, i.e. at the point of transfer as illustrated in **Figs. 4 and 7**. To determine the effect of learning and transfer on model parameters, we fit the data for each block of 50 trials using the collapsing bound model (without leak) described above.

None of the model parameters showed a dramatic change across trial blocks (**Fig. 10 CH**) except for drift rate, which increased gradually during training and then showed a sharper increase at the point of transfer from training to testing (**Fig. 10 A,B**, dashed vertical line). The drift rates were comparable for both eye movements and reaches, while the boundaries and collapse rates were somewhat greater for reaches (**Fig. 10, C-F**). Non-decision time (**Fig. 10 G,H**) was roughly 3 times longer for reaches than for eye movements.

**Figure 10.**
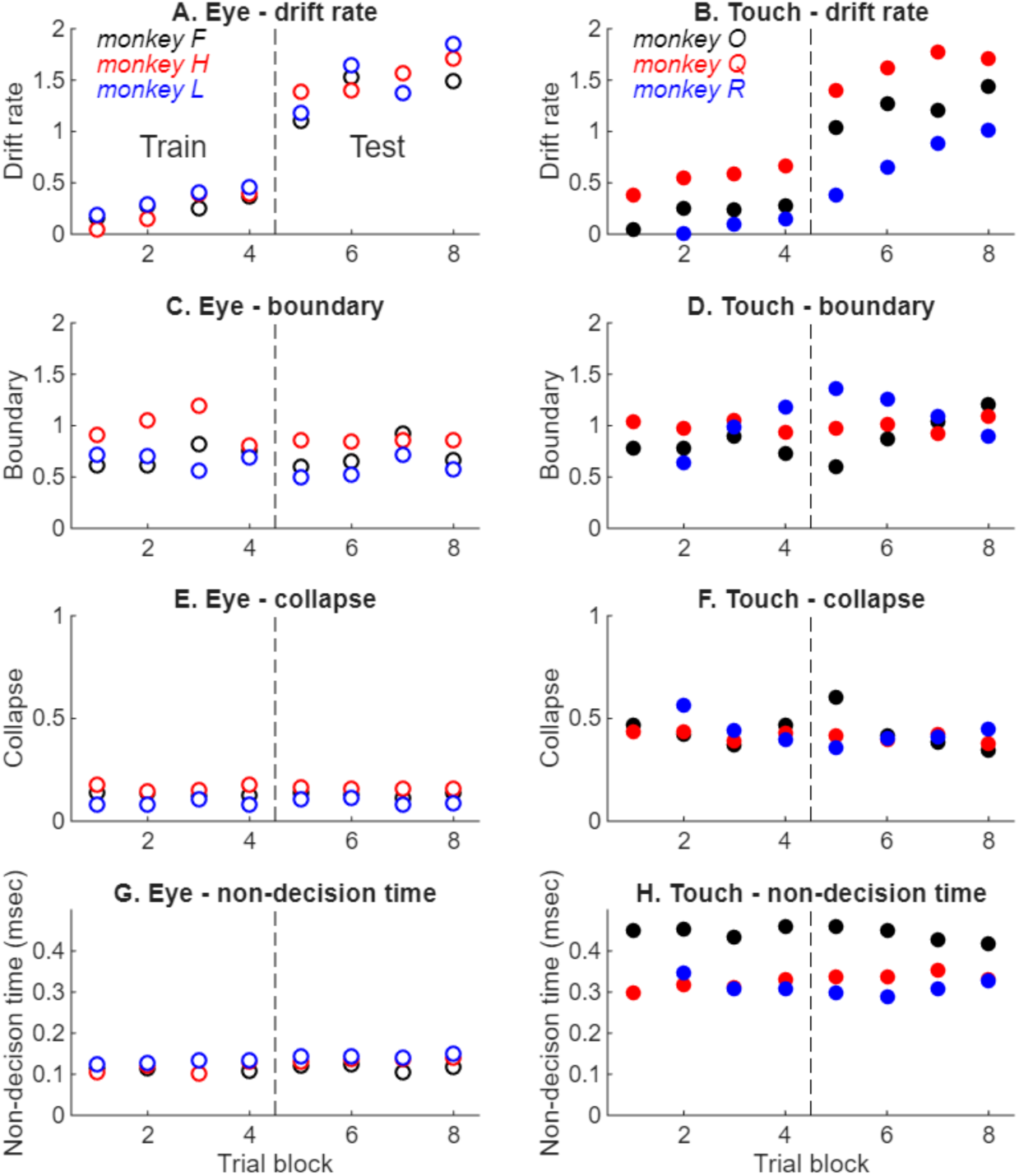
Parameter changes during learning and transfer. **A.** Drift rate for eye movement responses. **B.** Drift rate for reaching movements. **C.** Initial boundary position for eye movements. **D.** Boundaries for reaching movements. **E**. Collapse rate for eye movements. **F**. Collapse rate for reaches. **G**. Non-decision time for eye movements. **H**. Non-decision time for reaches. Dashed vertical lines indicate the point of transfer.

## Discussion

The transitive inference (TI) paradigm assesses the ability to infer the serial order of a set of stimuli based solely on pairwise comparisons, without explicit spatial or temporal cues. In this task, correct responses are rewarded for selecting the item of lower rank from a pair of images. When presented with adjacent pairs from a novel set of items, nonhuman primates (NHPs) initially performed at chance, but accuracy improved over the course of ~200-300 training trials. Accuracy then increased dramatically at the transition to all-pairs testing and showed a symbolic distance effect for non-terminal pairs, suggesting that subjects had acquired an internal representation of the list’s ordinal structure. How this internal structure and other latent variables contribute to decision making in this task was the subject of the current study.

To elucidate the mechanisms underlying TI decisions, we examined both accuracy and reaction time (RT) through the lens of evidence accumulation models. We found that choice RTs for eye movements were remarkably fast—generally under 300 milliseconds and only marginally longer (by 30–85 ms) than simple reaction times—suggesting that decisions were made rapidly even though there was little time pressure. Despite these rapid responses, performance improved reliably with training, implying that learning and emergence of an internal representation were supported even within this brief time window.

We initially explored whether a simple sequential sampling process could explain the pattern of results observed in the TI task. Simulations based on binomial likelihoods yielded an evidence accumulation rate highly sensitive to symbolic distance (**Fig. 8D**). However, this approach failed to capture the relative invariance of RT across SD conditions in the behavioral data (**Figs. 3D** and **4D**), motivating the need for a more flexible and biologically plausible framework.

The DDM provides a compelling theoretical framework for modeling decision dynamics. It allows for flexible manipulation of core parameters, including drift rate, decision boundary, and integration properties, resulting in distinct operational regimes. A constant bound model (**Fig. 2**, top row), which assumes a fixed decision threshold over time, has proven effective across many perceptual decision-making studies (Voskuilen et al., 2016). However, under conditions of low evidence (i.e., zero drift rate), this model predicts excessively long RTs unaccompanied by improvements in accuracy, a situation we refer to as the “zero-evidence catastrophe.”

To address this limitation, we explored models incorporating time-dependent (collapsing) boundaries (**Fig. 2**, bottom row). These models reduce the decision threshold over time, accelerating responses in low-evidence scenarios and yielding more symmetrical RT distributions—features more consistent with the observed behavioral data.

Another extension involved introducing leaky integration, where the accumulated evidence is affected by a decay (or amplification) factor. With a positive leak (unstable integration), accumulation becomes more rapid and boundary-crossing more likely, again minimizing long RTs (**Fig. 2**, middle row). However, fitting the current data using a constant-bound DDM required large positive leak values, suggesting that drift rate played a diminished role relative to the gain term, raising questions about the model’s plausibility. When leak was added to the collapsing bound model, it had a negligible or even deleterious effect on the fits. Importantly, the collapsing bound model without a leak term did not require extreme parameter values, making it a more parsimonious and biologically plausible account of TI decision-making.

In the TI transfer paradigm implemented in this study, learning was accompanied by a gradual increase in decision accuracy during training and a sharper increase at the point of transfer. This was found for both eye and reaching movements. The GDDM captured learning effects mainly through increases in drift rate, which reflect the increase in decision accuracy. There was little systematic change in other parameters with learning (**Fig. 10**).

As expected from previous studies, symbolic distance (SD)—the rank difference between paired stimuli—was a strong predictor of accuracy: larger SDs led to higher performance. However, the impact of symbolic distance on RT was minimal for both eye and reaching movements, consistent with prior work (Munoz et al., 2020). This dissociation between accuracy and RT contrasts with findings from perceptual decision-making paradigms, where RTs are typically longer (500–1000 ms) and inversely related to decision difficulty or accuracy (e.g., Roitman & Shadlen, 2002; Ratcliff et al., 2003; Kim & Shadlen, 1999; Kiani et al., 2014; Jun et al., 2021). In those studies, RT-accuracy trade-offs are commonly interpreted as signatures of gradual evidence accumulation, as formalized in sequential sampling models like the drift diffusion model (DDM).

Examining the empirical relationship between accuracy and RT (**Figs. 4** and **5**) revealed a weak but significant positive correlation during the early training phase. Within the DDM framework, this relationship reflects the interplay of drift rate and boundary height. Increases in drift rate reduce RT and improve accuracy, whereas increases in decision boundary prolong RT but also improve accuracy. The observed pattern suggests that, during learning, drift rate may have increased alongside modest adjustments in boundary height, enabling more accurate decisions at the cost of slightly delayed responses.

Reaction times for reaching responses were 2-3 times longer than eye movement responses. However, similar to eye movements, reach RTs showed little change over the course of learning. GDDM modeling revealed that the difference between eye movement and reach RTs was mainly attributed to non-decision time rather than a fundamental difference in decision dynamics. Previous studies of TI that used touchpanels (Gazes et al., 2012, 2014) reported significantly decreasing reaction times as a function of symbolic distance. These studies also reported changes in associative value that might be related to reward probability, which is directly related to decision accuracy and thus tends to increase with symbolic distance. This raises the possibility that reaction times might reflect motivational influences as well as decision processes. To mitigate the effects of reward associations, we used a different stimulus set in each behavioral session, we avoided pre-training on individual pairs, and we limited the overall length of each session. Nevertheless, the effects of motivation cannot be dismissed and deserve further investigation.

There are also studies indicating a dissociation between saccades reaction times and manual responses (Kveraga et al, 2002; Lawrence & Gardella 2009). In the current study, we did not test eye and reach movements in the same subject as this might have led to cross-training effects. Nevertheless, we found that eye and reach responses showed a similar dissociation between accuracy and reaction time albeit with substantially longer average response latencies for reaching movements. Future studies in which both response types are tested in the same subjects could reveal the degree to which they share the same or similar decision mechanisms.

The current findings provide new insights into the cognitive and computational processes underlying TI learning in nonhuman primates. By extending drift-diffusion modeling to a domain driven by internally inferred structure rather than direct sensory evidence, we highlight the adaptability of this decision framework to varying task demands. The success of the collapsing bound model suggests that decision policies are not fixed but evolve dynamically with learning, enabling fast yet accurate behavior even in abstract cognitive tasks.

This interpretation aligns with prior work suggesting that the utility of collapsing boundaries may depend on species and task context. While human data generally favor fixed-boundary models, with studies showing negligible practical advantages for collapsing variants (Voskuilen et al., 2016), evidence from nonhuman primates is more mixed. Some paradigms support collapsing boundaries, indicating that decision strategies may be more flexible in nonhuman species or under specific task demands. Hawkins et al. (2015) further emphasize that urgency signals and boundary dynamics can be modulated by contextual factors such as time pressure and task structure. In this light, our findings contribute to a growing body of evidence suggesting that decision boundaries are not static but can adapt to optimize performance in complex, internally guided tasks.

## Conclusion

The drift diffusion model (DDM) is traditionally employed in paradigms where accuracy and reaction time exhibit an inverse relationship. However, the flexibility of this modeling framework enables it to accommodate less conventional patterns of behavior, thereby offering insight into the interaction of latent cognitive variables across diverse decision-making contexts. In the present study, we demonstrate that a generalized DDM can be effectively applied to a reasoning task, transitive inference, where reaction times remain relatively constant despite significant learning-related improvements in accuracy. Our results suggest that, during the acquisition phase, the starting point of an exponentially collapsing decision boundary may be gradually modulated to reflect the evolving rate of evidence accumulation. This dynamic adjustment allows for an increased accuracy with minimal cost in response time, effectively enhancing decision efficiency. Furthermore, task performance was modulated by internal variables, such as symbolic distance, underscoring the role of internally computed metrics in guiding behavior. The collapsing boundary model provides a mechanistic explanation for these observations, minimizing the impact on reaction time as accuracy improves. In contrast, under conditions of slow evidence accumulation, this framework reduces the total evidence required by collapsing the decision threshold over time, acknowledging the diminishing returns of prolonged integration. Collectively, these findings highlight the utility of variable-bound DDMs in capturing the dynamics of decision-making under conditions where evidence is internally generated and temporally constrained.

## Data and code availability

The datasets and code generated and/or analyzed during the current study are available on Github.

## Acknowledgements

This study was supported by NIH R01-MH111703 (VPF and HST). No sponsor had any role in study design, data collection and analysis, decision to publish or preparation of the manuscript

## Author Contributions

The study was designed by FM, GJ, YA, VPF, and HST. Data were collected by FM and YA. Data were analyzed by FM, GJ, MS, JDM and VPF. The manuscript was written by FM, GJ, MS, and VPF.

